# SS18-SSX co-opts P300 to sustain oncogenic transcription independent of SWI/SNF activity

**DOI:** 10.64898/2026.01.26.701706

**Authors:** Anthony M. Doherty, Eimear Lagan, Richard J.R. Elliott, Keiran Wynne, Craig Monger, Adrian P. Bracken, Junwei Shi, Christopher R. Vakoc, Scott A. Armstrong, Giorgio Oliviero, Christopher Ott, Neil O. Carragher, Gerard L. Brien

## Abstract

Synovial sarcoma is driven by the SS18-SSX fusion oncoprotein, which has been assumed to promote tumorigenesis through its incorporation into the SWI/SNF chromatin remodeling complexes. Accordingly, therapeutic efforts have focused on targeting SS18-SSX containing SWI/SNF assemblies, yet these approaches have produced limited clinical benefit. Here, we demonstrate that SS18-SSX sustains oncogenic transcription independent of SWI/SNF activity. Despite efficient degradation and dismantling of SWI/SNF complexes, fusion occupancy at target loci and associated gene expression programs remain largely intact. Instead, we identify the acetyltransferase P300 as an essential co-factor supporting SS18-SSX chromatin binding and transcriptional activation. Targeting P300 displaces the fusion from chromatin, suppresses its transcriptional output, compromising synovial sarcoma viability. Notably, dual PROTAC mediated degradation of P300 and SWI/SNF produces strong synergistic effects, broadly disrupting SS18-SSX localization and function. These findings redefine the mechanistic basis of synovial sarcoma and reveal a mechanistically anchored therapeutic strategy for targeting its core oncogenic driver.

## INTRODUCTION

Synovial sarcoma is an aggressive soft-tissue malignancy primarily effecting children and young adults^1^. Disease development is driven by a pathognomonic chromosomal rearrangement present in essentially 100% of patients t(X;18) (p11.2;q11.2), leading to the generation of the SS18-SSX oncogenic fusion protein^2-4^. Sarcoma genomics studies have found limited evidence of any additional highly recurrent genetic changes in these tumours^5-7^. Moreover, modelling the disease in mice shows that SS18-SSX is sufficient to drive tumour development^8,9^. Therefore, it is clear that the SS18-SSX fusion protein is the core disease driver. As such, discovering the oncogenic mechanisms utilised by the fusion is essential, both for our understanding of disease biology and to guide the development and implementation of disease-centred therapeutics. Advances in this regard are much needed, since the long-term survival rate for synovial sarcoma patients remains ∼50%, with no mechanistically anchored therapeutics currently in clinical use^1,10,11^.

SS18 is a component of the mammalian SWI/SNF chromatin remodelling complexes, while the SSX component, which can be any one of 3 closely related genes (SSX1, SSX2 or SSX4) appears to function as a transcriptional repressor^2-4,12-14^. SWI/SNF complexes are modular, multi-protein complexes that assemble into 3 distinct molecular subclasses – canonical BAF (cBAF), non-canonical BAF (ncBAF, also known as GLTSCR-associated BAF (GBAF)) and Polybromo-associated BAF (PBAF) (Figure S1A)^15-18^. SS18-SSX dominantly assembles into 2 of these subclasses (cBAF and ncBAF), leading to the redistribution of fusion containing complexes on chromatin^12,13,19,20^. As such, this fusion event is thought to create a potent epigenetic driver that reprograms gene expression to drive and sustain tumour development. It is thought that the fusion primarily exploits the chromatin remodelling activity of the SWI/SNF complexes into which it incorporates, to drive oncogenic gene expression. Indeed, supporting this we and others have identified functional dependencies on ncBAF complexes in synovial sarcoma cells, particularly on the BRD9 component of these complexes^16,21^. This motivated our development of a Proteolysis Targeting Chimera (PROTAC)-based approach to degrade BRD9 in cells^21,22^. Despite promising preclinical results and robust degradation activity in tumours, BRD9 PROTACs have provided only transient patient responses^21,23^. This highlights the need for a more detailed understanding of the oncogenic mechanisms utilised by SS18-SSX, to develop alternative and/or complimentary therapeutic approaches for this disease.

Here, we demonstrate that SS18-SSX supports oncogenic gene expression independent of SWI/SNF activity. PROTAC mediated targeting of SWI/SNF leading to the destruction of fusion containing complexes has limited impact on SS18-SSX associated gene expression. Instead, we found that the fusion co-opts the activity of P300 to support oncogenic gene expression. Therapeutic targeting of P300 displaces SS18-SSX from chromatin, diminishing associated gene expression programs. Remarkably, we found combined PROTAC treatment targeting SWI/SNF and P300, was highly synergistic in synovial sarcoma cells. Combination treatments profoundly disrupted SS18-SSX localisation, associated gene expression programs and sarcoma cell viability. Therefore, we have discovered a mechanistically anchored therapeutic strategy disrupting the core disease driver of synovial sarcoma tumour biology.

## RESULTS

### SWI/SNF complex dynamics following BRD9 degradation

We previously demonstrated that degradation of BRD9 has therapeutic potential in synovial sarcoma, but clinical testing has yielded transient responses^21,23^. To define how BRD9 degradation impacts SS18-SSX and SWI/SNF complexes in synovial sarcoma, we established an endogenous purification platform to analyse SWI/SNF assemblies (**Figure 1A** and **S1A**). Specifically, we purified complexes via the shared core subunit SMARCC1, present in all SWI/SNF subtypes; and in parallel via SS18-SSX to directly interrogate fusion protein containing complexes (**Figure 1A**). These purifications robustly enriched SWI/SNF components, enabling detailed analyses of complex assembly in cells. Consistent with recent work, PBAF appeared to be the most abundant SWI/SNF subclass in synovial sarcoma cells (**Figure S1B**)^24^. However, direct purification of SS18-SSX revealed that the majority of fusion containing complexes (∼65%) were the cBAF subclass, with the remainder corresponding to BRD9 containing ncBAF (**Figure S1B-D**). As expected, given that SS18-SSX is not a PBAF component, we observed only marginal enrichment of PBAF-specific components in SS18-SSX purifications (**Figure S1B-D**). Together, these data establish a robust system to examine SWI/SNF dynamics in synovial sarcoma.

**Figure 1:**
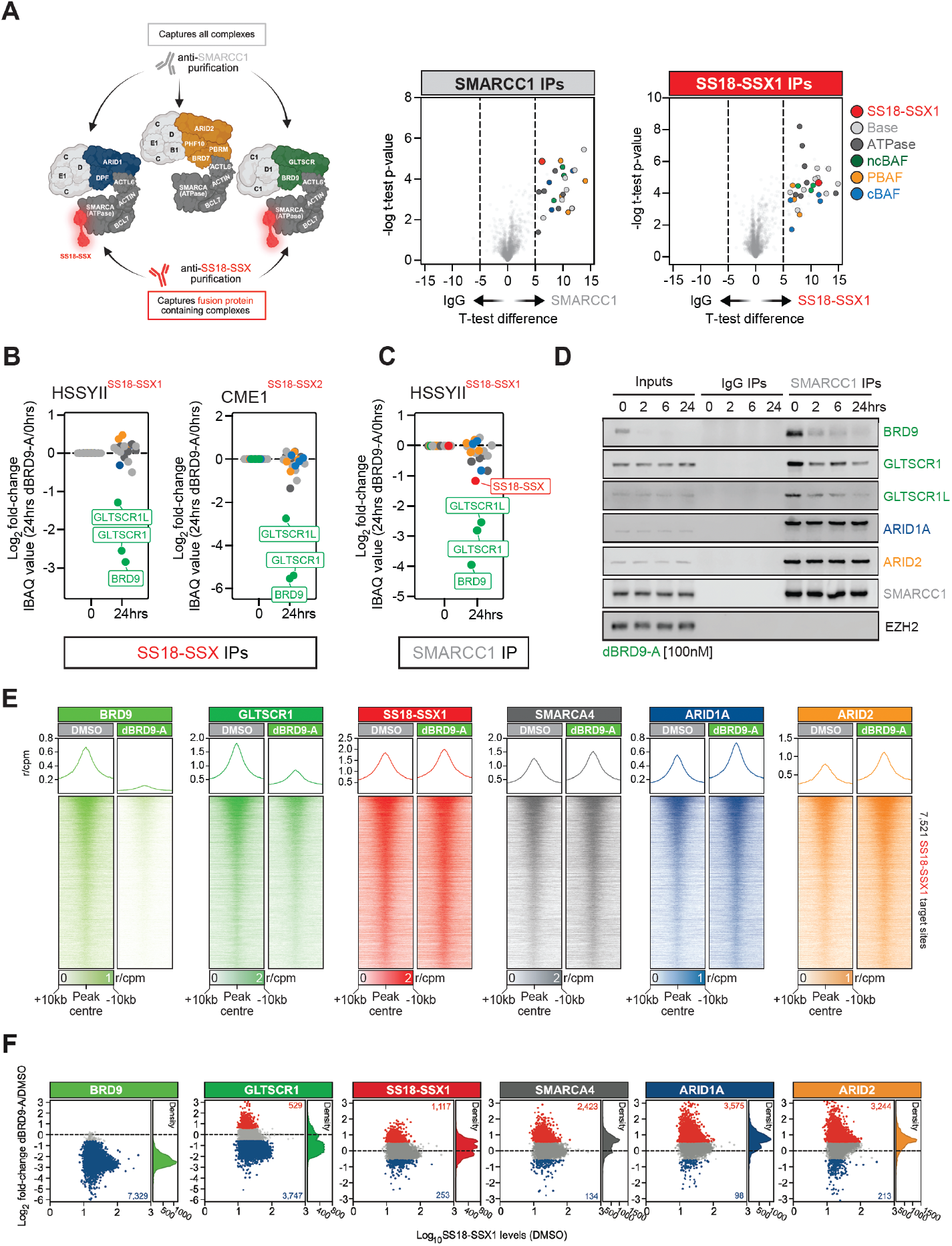
SWI/SNF and SS18-SSX dynamics in BRD9 degrader treated cells. **A**. Schematic of SWI/SNF complex purifications using SMARCC1 and SS18-SSX antibodies (left panel). Volcano plots of SMARCC1 and SS18-SSX purification mass spec data from HSSYII cells. SWI/SNF complex members are colour coded and indicated on the plot. **B**. Abundance plots depicting SWI/SNF member fold-change in SS18-SSX purifications following 24hrs of dBRD9-A 100nM treatment in HSSYII and CME1 cells. SWI/SNF members are coloured as per panel A. **C**. Abundance plots depicting SWI/SNF member fold-change in SMARCC1 purifications following 24hrs of dBRD9-A 100nM treatment in HSSYII cells. **D**. Immunoblots of the indicated SWI/SNF members and EZH2 in SMARCC1 purifications in HSSYII cells treated with dBRD9-A 100nM for 2, 6 and 24hrs. **E**. Tornado and average plots showing the abundance of SWI/SNF members in quantitative CUT&RUN-Rx of control (DMSO) and dBRD9-A 100nM treated (24hrs) HSSYII. Scale denotes reference-adjusted counts per million mapped reads (r/CPM). **F**. MA plots showing changes in SWI/SNF member abundance at SS18-SSX target sites following 24hrs of dBRD9-A 100nM treatment. Sites changing >1.5-fold are shown in red (increasing) or blue (decreasing) with the number of sites in these categories indicated.

We next asked how BRD9 degradation alters SWI/SNF complex assembly. SS18-SSX purifications were performed in two independent fusion positive cell lines, HSSYII (SS18-SSX1) and CME1 (SS18-SSX2) (**Figure 1B**). BRD9 degradation produced a highly selective impact, the only proteins significantly depleted from SS18-SSX purifications were BRD9 and the ncBAF defining subunits GLTSCR1 and GLTSCR1L16,17, which were all lost from fusion containing complexes (**Figure 1B**). Since SS18-SSX purifications do not capture PBAF, we extended these analyses to include SMARCC1 purifications. As in SS18-SSX purifications, ncBAF defining members were selectively lost upon BRD9 degradation with few additional changes (**Figure 1B-C**). Notably, the next most affected protein was SS18-SSX itself, which was reduced in SMARCC1 purifications following BRD9 loss (**Figure 1C**). These data indicate BRD9 degradation leads to dissolution of ncBAF complexes, including fusion containing ncBAF, while other SWI/SNF assemblies remain largely intact (**Figure 1B-D** and **S1B-D**). The persistence of these complexes provides a potential mechanistic basis for the transient clinical responses to BRD9 degraders.

### SWI/SNF chromatin binding dynamics following BRD9 degradation

To determine whether the remaining complexes functionally compensate for ncBAF loss, we mapped their chromatin occupancy following BRD9 degradation. We used a quantitative CUT&RUN strategy with exogenous genome spike-in normalisation to profile SWI/SNF members in PROTAC treated cells^25^. As expected, BRD9 signal was broadly depleted across SS18-SSX bound regions (**Figure 1E-F** and **S1E**). Chromatin bound levels of GLTSCR1 were also markedly reduced, with ∼50% of sites exhibiting >1.5-fold loss of GLTSCR1 (**Figure 1E-F** and **S1E**). A subset of loci retained or modestly gained GLTSCR1 binding, suggesting that GLTSCR1 can associate with chromatin at some regions independently of fully assembled ncBAF (**Figure 1E-F**).

We then examined how these changes affect SS18-SSX and the remaining SWI/SNF subclasses. Despite reduced incorporation of SS18-SSX in ncBAF complexes (**Figure 1C**), fusion occupancy was largely maintained after BRD9 degradation (**Figure 1E**). Global SS18-SSX signal was modestly increased across its target regions, with ∼15% of sites showing >1.5-fold gains in fusion binding, although a minority of loci exhibited decreased occupancy (**Figure 1E-F** and **S1E**). Profiling the shared ATPase subunit SMARCA4, together with cBAF specific ARID1A and PBAF specific ARID2 revealed increased levels of these factors at fusion bound regions following BRD9 degradation (**Figure 1E** and **S1E-F**). Between 30-50% of fusion bound loci gained >1.5-fold binding of these proteins (**Figure 1F** and **S1F**). Increases in SS18-SSX correlated with gains in SMARCA4 and ARID1A, but not ARID2 (**Figure S1G**); indicating that SS18-SSX containing cBAF complexes increase in the absence of ncBAF, while PBAF recruitment is largely uncoupled from the fusion. Collectively, these data show that BRD9 degradation triggers a redistribution of SWI/SNF activity, with increased engagement of remaining complexes at SS18-SSX target sites (**Figure S1H**).

### Broader therapeutic disruption of SS18-SSX containing SWI/SNF complexes

We next tested whether a broader disruption of SS18-SSX containing complexes could mitigate any potential compensatory activity. Quantitative analysis of fusion purifications identified SMARCA4 and ACTL6A as the most abundant SWI/SNF members associated with SS18-SSX (**Figure S2A-B**). This is consistent with their presence, with wildtype SS18 within the ATPase module of SWI/SNF^15,26,27^; and indicates that SS18-SSX retains a similar architecture within the complex. Importantly, these data show the vast majority of SS18-SSX is stably associated with SMARCA4, or its paralog SMARCA2. Highlighting these proteins as therapeutic targets with the broadest potential to disrupt fusion containing complexes (**Figure S2A-B**).

Recent PROTAC development has provided small molecules targeting SMARCA2/4^28-30^. Treatment of synovial sarcoma cells with ACBI-1, potently degrades SMARCA2 and SMARCA4, with reduced activity against the PBAF component PBRM1, led to extensive disassembly of SWI/SNF complexes (**Figure 2A**). SMARCC1 purifications showed that the SWI/SNF core module remained intact and retained interactions with cBAF subunits. However, all PBAF, ncBAF and ATPase module components were lost, indicating that no fully assembled SWI/SNF complexes remained (**Figure 2A**). SS18-SSX purifications under these conditions revealed complete dissolution of fusion containing complexes. Core, ncBAF and ATPase subunits were all lost from SS18-SSX pulldowns in ACBI-1 treated cells (**Figure 2A**). Thus, SMARCA2/4 degradation ablates assembly of SS18-SSX containing SWI/SNF complexes.

**Figure 2:**
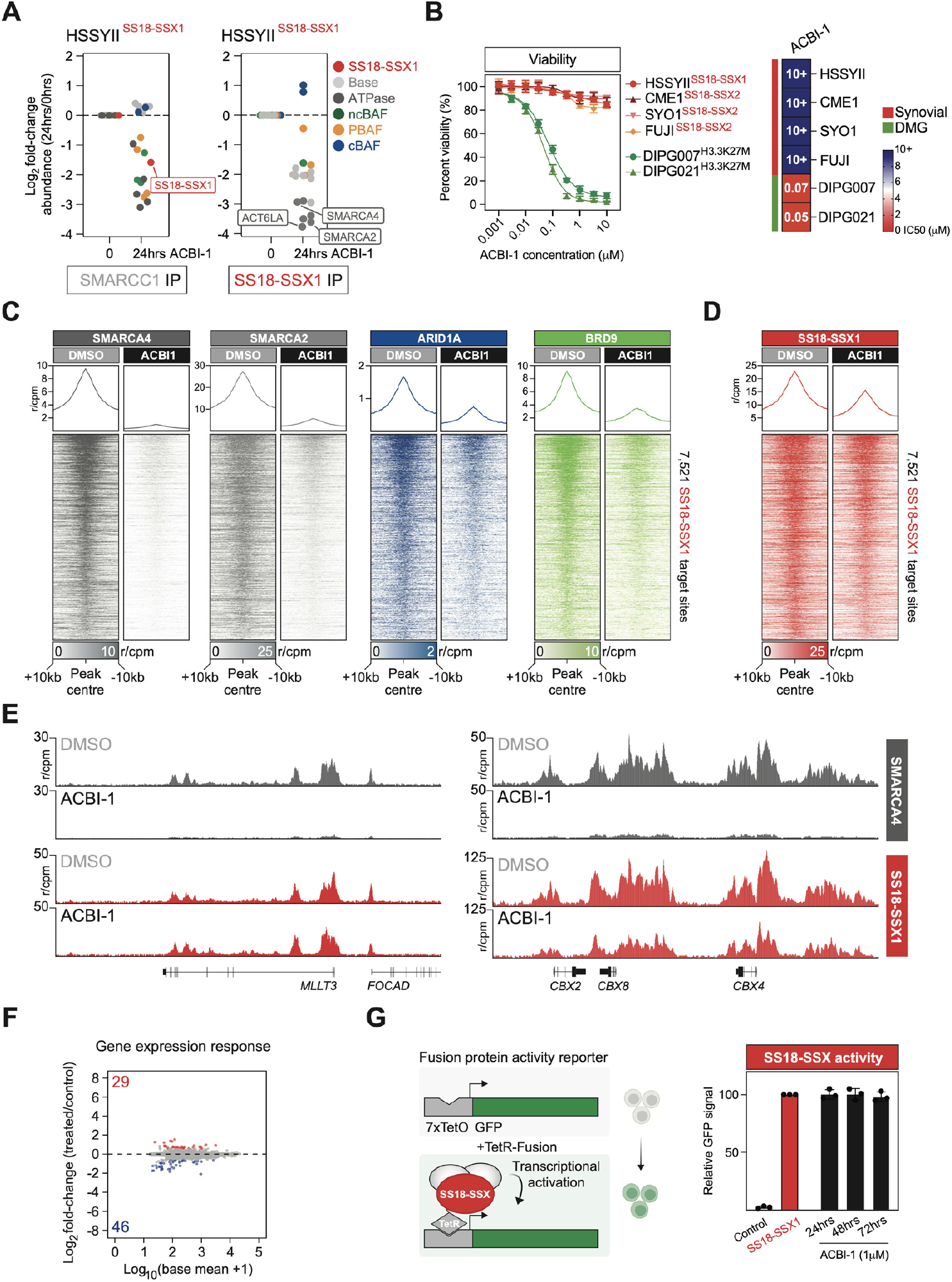
SWI/SNF complexes are dispensable for SS18-SSX mediated transcription. **A**. Abundance plots depicting SWI/SNF member fold-change in SS18-SSX and SMARCC1 purifications following 24hrs of 1μM ACBI-1 treatment in HSSYII cells. SWI/SNF members are colour coded as shown. **B**. Cell viability dose response data in the indicated synovial sarcoma and diffuse midline glioma cell lines treated with ACBI-1 for 72hrs. Mean +/-s.d., n = 3 (left panel). Heatmap representing ACBI-1 IC50 doses in each cell line examined. **C**. Tornado and average plots showing the abundance of the indicated SWI/SNF members in CUT&RUN-Rx of control (DMSO) and 24hrs 1μM ACBI-1 treated HSSYII cells. Scale denotes reference-adjusted counts per million mapped reads (r/CPM). **D**. Tornado and average plots showing abundance of SS18-SSX as per panel C. **E**. Genomic tracks showing SS18-SSX and SMARCA4 signal at the indicated genomic loci in control (DMSO) and ACBI-1 treated HSSYII cells. **F**. MA plot showing differential gene expression in HSSYII cells following 24hrs of 1μM ACBI-1 treatment. Significantly differentially expressed genes are indicated in red (upregulated) and blue (downregulated). **G**. Schematic showing the GFP transcriptional reporter system (left panel). Bar plot showing changes in GFP reporter gene expression in TetR-SS18-SSX expressing cells, treated with ACBI-1 at the indicated time points.

### SWI/SNF complexes are not required for oncogenic gene expression

Despite the near complete dismantling of SWI/SNF complexes, ACBI-1 treatment had minimal short-term impact on synovial sarcoma cell viability in dose response assays (**Figure 2B**). In contrast, H3K27M-mutant diffuse midline glioma cells, which depend on SMARCA4 function showed pronounced sensitivity to ACBI-1 (**Figure 2B**)^29,31^. This suggests synovial sarcoma cells lack an acute dependency on SWI/SNF function.

To further define the consequences of SMARCA2/4 degradation, we performed quantitative CUT&RUN profiling of multiple SWI/SNF components (**Figure 2C-D** and **S2C**). As expected, SMARCA2 and SMARCA4 occupancy was strongly reduced at SS18-SSX target sites in ACBI-1 treated cells (**Figure 2C** and **S2C**). Consistent with the broad disruption of complex assembly, ARID1A and BRD9 binding was also markedly reduced, indicating loss of cBAF and ncBAF at these loci (**Figure 2C**). Approximately, 85-90% of SS18-SSX bound regions exhibited >1.5-fold reductions in ARID1A and BRD9 (**Figure S2C**), confirming that SMARCA2/4 degradation broadly compromises SS18-SSX containing cBAF and ncBAF complexes.

Mapping SS18-SSX in these conditions revealed an overall decrease in fusion occupancy at target loci (**Figure 2D**). However, the magnitude of this reduction was modest relative to ARID1A and BRD9; with ∼50% of sites showing >1.5-fold loss of SS18-SSX signal. Moreover, many loci retained high levels of fusion protein binding despite loss of the associated complex subunits (**Figure 2E** and **S2C**). The SSX domain of the fusion can directly interact with chromatin, particularly H2AK119ub modified domains^32,33^. The persistence of SS18-SSX at target sites in the absence of intact SWI/SNF complexes suggests that this intrinsic chromatin binding activity and/or additional mechanisms support fusion occupancy independent of SWI/SNF. DepMap analyses indicate that genetic disruption of SMARCA4 has a measurable phenotypic impact in synovial sarcoma cells (**Figure S2D**); and our independent genetic perturbation experiments similarly show that long-term targeting of SMARCA2/4 reduces viability (**Figure S2E)**. Moreover, extended ACBI-1 treatment can eventually elicit a viability response in SS18-SSX positive lines (**Figure S2F**). The requirement for prolonged SMARCA2/4 suppression prior to observing any phenotypic effect, suggests that although SWI/SNF complex activity is required for long-term viability, it is not directly responsible for maintaining oncogenic transcriptional output.

To test this, we mapped global gene expression in ACBI-1 treated synovial sarcoma cells. Remarkably, despite the broad disruption of SWI/SNF complexes, transcriptional changes were minimal (**Figure 2F**). We then used a reporter system to directly assess SS18-SSX driven transcription (**Figure 2G**). Recruitment of SS18-SSX to a TetO-GFP locus robustly activated the reporter. However, ACBI-1 treatment did not diminish reporter expression (**Figure 2G**). Thus, SWI/SNF activity is dispensable for SS18-SSX mediated transcription. Collectively, these results suggest that the limited clinical efficacy of SWI/SNF targeting in synovial sarcoma reflects a SWI/SNF independent mechanism employed by SS18-SSX to drive oncogenic transcriptional programs.

### P300 is a functionally essential SS18-SSX co-factor

We next sought to identify chromatin regulators required for oncogenic transcription. We performed a focus CRISPSR/Cas9 screen targeting functional domains in ∼250 chromatin regulators (**Figure 3A**)^21,34^. This screen revealed a specific dependency on the acetyltransferase and bromodomain regions of P300 (**Figure 3A**). Notably, this effect was restricted to P300 and not its paralog CBP, despite comparable expression of both genes (**Figure S3A** and data not shown). Parallel experiments in EWS-FLI1 positive Ewing sarcoma cells showed no similar dependency, indicating that the P300 dependence is specific to synovial sarcoma cells (**Figure 3A** and **S3A-B**). Independent DepMap analyses supported this, revealing a strong correlation between SS18 and P300 dependency in synovial sarcoma cell lines (r = 0.757), whereas other cancer types showed no such relationship (r = 0.135) (**Figure 3B**). Consistent with this being linked to the presence of the fusion, the SW982 cell line, designated as synovial sarcoma but lacking SS18-SSX showed no P300 dependency (**Figure 3B**). Together, these data demonstrate that SS18-SSX positive synovial sarcoma cells are functionally reliant on P300, implicating P300 as a key SS18-SSX co-factor.

**Figure 3:**
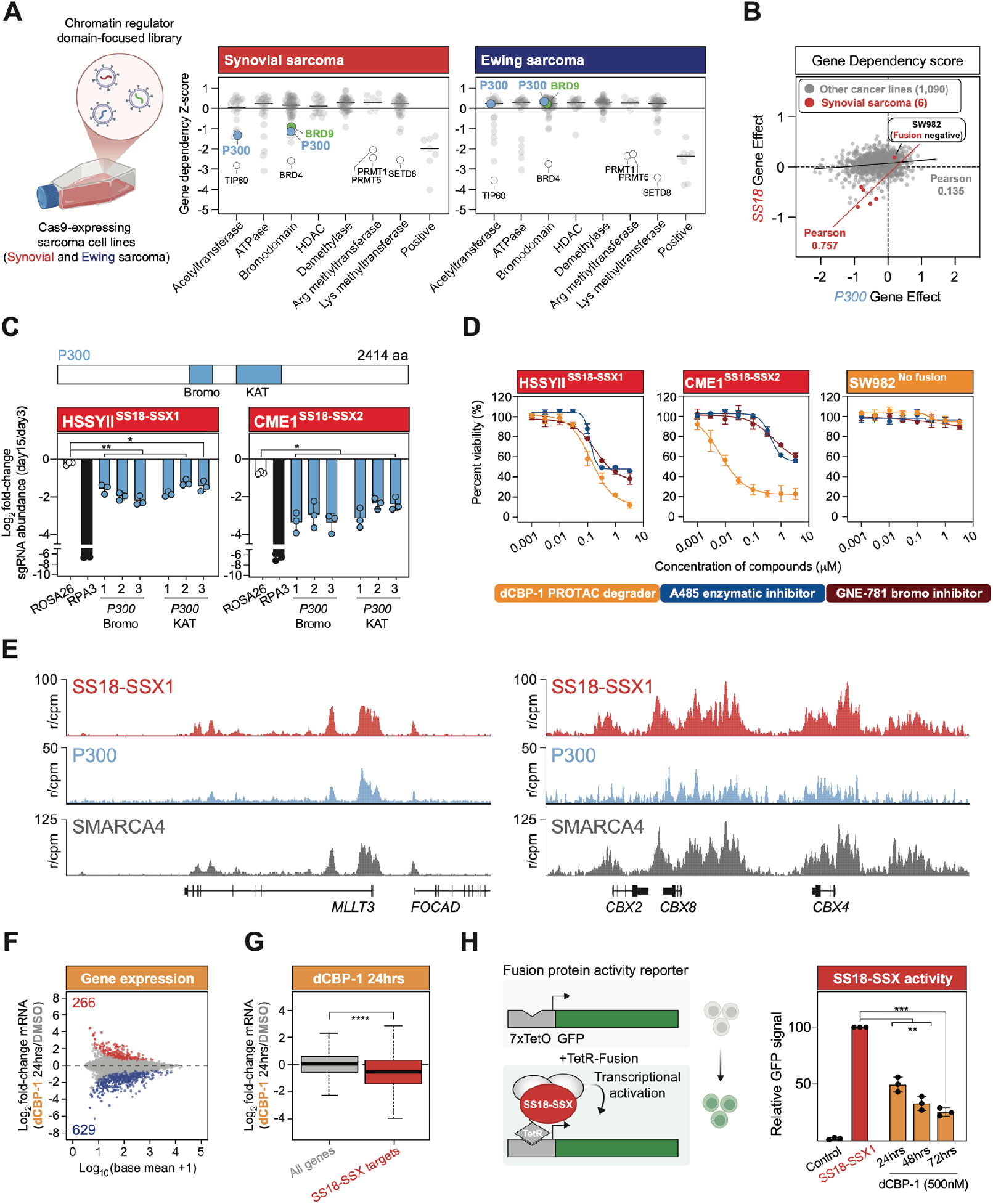
P300 is a functionally essential SS18-SSX transcriptional co-factor. **A**. CRISPR screening gene dependency Z-Scores for sgRNAs targeting functional domains in chromatin regulatory genes. Screens were performed in multiple synovial and Ewing sarcoma cell lines; each dot denotes average Z-Score for domain targeting sgRNAs across cell lines. **B**. Dot plot showing CRISPR gene effect scores for SS18 and P300 in cell lines within DepMap. Synovial sarcoma cell lines are highlighted in red. Pearson correlations are shown for synovial (red) and all other (grey) cell lines. **C**. Growth competition assays in the indicated synovial sarcoma lines expressing three independent P300 bromodomain or acetyltransferase targeting sgRNAs (n = 3, data represents mean +/-SD). P-values calculated using Student’s t-test comparing against negative control sgRNA targeting ROSA26, * = P ≤ 0.05, ** = P ≤ 0.01. **D**. Cell viability dose response data in the indicated synovial sarcoma cell lines treated with P300/CBP targeting small-molecules (inhibitors (A-485 and GNE-781) and degrader (dCBP-1)). Mean +/-s.d., n = 3 (left panel). **E**. Genomic tracks showing quantitative CUT&RUN-Rx signal for SS18-SSX, P300 and SMARCA4 at the indicated genomic loci in control HSSYII cells. **F**. MA plot showing differential gene expression in HSSYII cells following 24hrs of dCBP-1 500nM treatment. Significantly differentially expressed genes are indicated in red (upregulated) and blue (downregulated). **G**. Boxplot showing Log2 fold-change of direct SS18-SSX target genes (red) and all other genes (grey) in HSSYII cells following 24hrs of dCBP-1 500nM treatment. **H**. Schematic showing the GFP transcriptional reporter system (left panel). Bar plot showing changes in GFP reporter gene expression in TetR-SS18-SSX expressing cells, treated with 500nM dCBP-1 at the indicated time points.

The acetyltransferase and bromodomain of P300 have proven to be tractable drug targets^35-44^. sgRNAs targeting either domain produced strong and specific, viability defects in synovial sarcoma cell lines (**Figure 3C** and **S3B**). In line with this, small-molecule inhibition of the P300 acetyltransferase domain (A-485) or bromodomain (GNE-781) elicited a marked viability response in SS18-SSX positive cell lines, while the fusion negative SW982 cell line was insensitive to both inhibitors (**Figure 3D** and **S3C-D**). Since both domains are functionally required, we reasoned that targeted degradation might provide greater efficacy compared to inhibitor treatment. Indeed, SS18-SSX positive lines were highly sensitive to the P300/CBP PROTAC dCBP-141, which produced stronger therapeutic responses than catalytic or bromodomain inhibitors, while SW982 cells remained insensitive (**Figure 3D** and **S3E**).

Wildtype SS18 has been reported to interact with P300^45^. We therefore wondered whether SS18-SSX retains this interaction. SS18-SSX purifications from multiple cell lines revealed co-purification of P300 and CBP, demonstrating that the fusion is physically associated with P300 in sarcoma cells (**Figure S3F**). Consistent with this interaction, mapping P300 localisation on chromatin revealed extensive co-localisation with SS18-SSX (**Figure 3E** and **S3G**). Ranking all fusion bound regions into deciles based on SS18-SSX abundance demonstrated that P300 occupancy closely tracked with fusion binding, with the strongest SS18-SSX target sites coinciding with the most highly P300 bound and H3K27ac marked regions (**Figure S3H**). Since SS18-SSX has been proposed to bind chromatin in part via interactions with PRC1-mediated H2AK119ub, we also examined Polycomb-associated marks at fusion bound sites. H3K27me3 and H2AK119ub were enriched at SS18-SSX bound sites (**Figure S3G-H)**. However, regions with high P300 levels exhibited markedly reduced Polycomb marks, consistent with the antagonistic relationship between P300 mediated acetylation and Polycomb complex function^46,47^. These observations suggest SS18-SSX may exploit alternative recruitment mechanisms at highly P300 bound sites, independent from established mechanisms.

### P300 supports SS18-SSX driven transcriptional output

To define the role of P300 in transcriptional control, we treated synovial sarcoma cells with dCBP-1, performing global expression analyses. In contrast to ACBI-1, dCBP-1 induced a broad transcriptional response (**Figure 3F**). Notably, this response included robust downregulation of direct fusion protein target genes, indicating that P300 is essential for maintaining SS18-SSX driven transcription (**Figure 3G**). We examined this further using the SS18-SSX transcriptional reporter. dCBP-1 treatment markedly reduced reporter expression (**Figure 3H**); demonstrating that P300 activity is required for SS18-SSX mediated transcriptional activation. Collectively, these data show P300 directly supports the oncogenic transcriptional output associated with SS18-SSX.

### P300 supports SS18-SSX chromatin binding

Given that P300 is required for SS18-SSX driven transcription, we next examined whether P300 also contributes to fusion protein localisation on chromatin. We performed quantitative CUT&RUN profiling of SS18-SSX, P300 and CBP in dCBP-1 treated cells (**Figure 4A** and **S4A**). Degradation of P300/CBP resulted in a pronounced reduction in SS18-SSX occupancy at its target sites. Reductions in P300 (and CBP) closely correlated with decreases in SS18-SSX (**Figure 4B** and **S4B**); indicating that P300 directly promotes fusion protein binding. To examine this relationship more systematically, we stratified fusion bound regions according to P300 abundance (**Figure 4C**). Regions with the highest P300 signal showed the strongest loss of SS18-SSX upon dCBP-1 treatment, whereas sites with low P300 levels were affected to a much lesser extent (**Figure 4C-D**). Strikingly, the overall reduction in SS18-SSX binding following P300 degradation was substantially greater than that observed with ACBI-1. Approximately 70% of sites lost >1.5-fold SS18-SSX signal after dCBP-1 treatment (**Figure S4C**), compared with only ∼50% with SMARCA2/4 degradation (**Figure S2C**). Thus, these results demonstrate that P300 is required to support SS18-SSX chromatin binding at their shared target sites. Since ACBI-1 treatment also decreased SS18-SSX binding, albeit more modestly, we compared the chromatin binding defects induced by ACBI-1 and dCBP-1. Remarkably, ACBI-1 primarily displaced the fusion from sites with low P300 levels, regions that are largely insensitive to dCBP-1 treatment (**Figure 4C&E-F**). Therefore, ACBI-1 and dCBP-1 have distinct impacts on SS18-SSX localisation.

**Figure 4:**
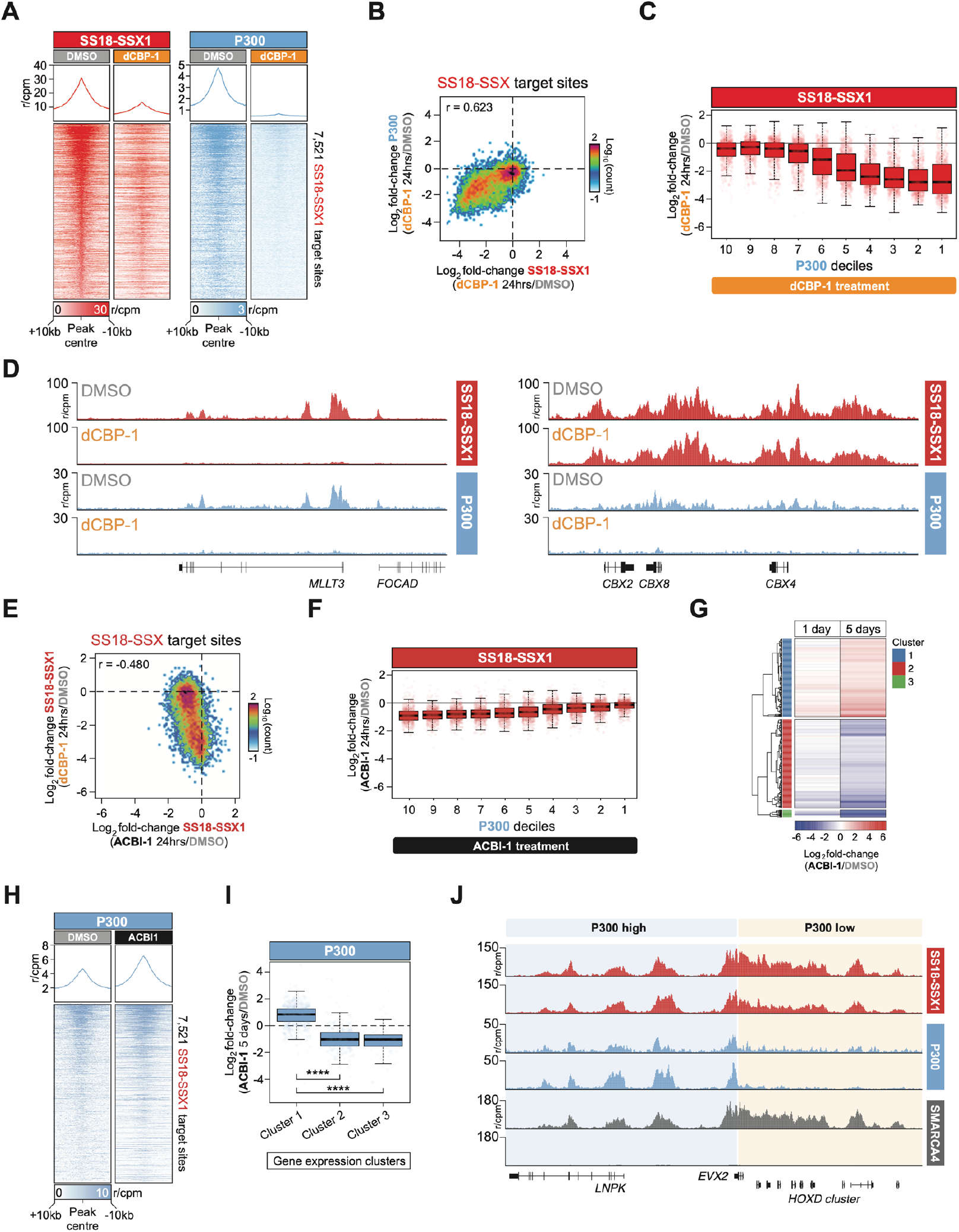
P300 is required for SS18-SSX chromatin binding. **A**. Tornado and average plots showing the abundance of SS18-SSX and P300 in CUT&RUN-Rx of control (DMSO) and 24hrs dCBP-1 500nM treated HSSYII cells. Scale denotes reference-adjusted counts per million mapped reads (r/CPM). **B**. XY scatter plot correlating changes in P300 and SS18-SSX CUT&RUN-Rx signal at fusion protein target sites in 24hrs dCBP-1 500nM treated HSSYII cells. **C**. P300 decile boxplots showing Log2 fold-change in SS18-SSX CUT&RUN-Rx signal at fusion protein target sites in 24hrs dCBP-1 500nM treated HSSYII cells. **D**. Genomic tracks showing CUT&RUN-Rx signal for SS18-SSX and P300 at the indicated genomic loci in control (DMSO) and 24hrs dCBP-1 500nM treated HSSYII cells. **E**. XY scatter plot correlating changes in SS18-SSX CUT&RUN-Rx signal at fusion protein target sites in 24hrs dCBP-1 (Y-axis) and ACBI-1 (X-axis) treated HSSYII cells. **F**. P300 decile boxplots showing Log2 fold-change in SS18-SSX CUT&RUN-Rx signal at fusion protein target sites in 24hrs ACBI-1 1μM treated HSSYII cells. **G**. Heatmap showing Log2 fold-change in mRNA abundance of differentially expressed direct SS18-SSX target genes in 5-day ACBI-1 1μM treated cells. **H**. Tornado and average plots showing the abundance of P300 in CUT&RUN-Rx of control (DMSO) and 5-day ACBI-1 1μM treated HSSYII cells. Scale denotes reference-adjusted counts per million mapped reads (r/CPM). **I**. Boxplots showing Log2 fold-change in P300 CUT&RUN-Rx signal at peaks associated with differentially expressed SS18-SSX target gene clusters in panel G. **J**. Genomic tracks showing SS18-SSX, P300 and SMARCA4 CUT&RUN-Rx signal at the indicated genomic loci in control (DMSO) and 5-day ACBI-1 1μM treated HSSYII cells. Region is divided based on high and low levels of P300 binding at fusion target sites.

We had seen that prolonged SMARCA2/4 degradation eventually reduces synovial sarcoma viability (**Figure S2F**). We therefore asked whether extended ACBI-1 treatment leads to greater displacement of SS18-SSX from chromatin. However, quantitative mapping of SMARCA4 and SS18-SSX after 5 days of ACBI-1 treatment revealed similar reductions in SS18-SSX binding (**Figure S4D**), compared to those observed at 24hrs (**Figure 2C-D** and **S4D**). Longer term ACBI-1 treatment elicited a broader transcriptional response compared with short-term treatment (**Figure 2F** and **S4E**). Although this was not associated with global downregulation of SS18-SSX target genes (**Figure S4F**). Instead, differentially expressed genes included comparable numbers of up- and downregulated SS18-SSX target genes (**Figure 4G**). Given that P300 plays a central role in fusion driven transcription, we mapped P300 localisation following 5-day ACBI-1 treatment. Notably, P300 levels increased significantly at fusion bound sites (**Figure 4H**). We examined P300 dynamics specifically at differentially expressed fusion protein target genes (**Figure 4G**). Remarkably, P300 occupancy increased at upregulated SS18-SSX target genes and decreased at downregulated genes (**Figure 4I**), suggesting that changes in P300 binding are linked to the observed transcriptional outcomes. Consistent with this, upregulated genes displayed higher P300 levels in untreated cells and P300 dynamics correlated with SS18-SSX changes; with increasing P300 abundance being associated with sustained or enhanced SS18-SSX binding (**Figure 4J** and **S4H-J**). These results demonstrate P300 supports SS18-SSX binding and transcriptional output independent of SWI/SNF, while sites with low P300 remain susceptible to SWI/SNF disruption.

### Synergistic therapeutic impact of combined degrader treatment

Since ACBI-1 and dCBP-1 primarily displace SS18-SSX from distinct subsets of target sites, and because P300 sustains fusion protein binding in ACBI-1 treated cells, we reasoned that simultaneous degradation of SMARCA2/4 and P300/CBP might produce synergistic therapeutic effects. To test this, we performed dose-response experiments using a 7×7 dose-ration matrix and formal synergy analysis to compare each degrader alone and in combination. Remarkably, combined ACBI-1 and dCBP-1 treatment produced strong synergy across all SS18-SSX positive cell lines tested (**Figure 5A** and **S5A**), whereas no significant synergy was observed in the fusion negative SW982 cell line (**Figure 5A**). The magnitude of this synergy enabled profound therapeutic responses even at substantially reduced concentrations of each individual compound when used in combination (**Figure 5A** and **S5A-B**). This synergy suggests that dual targeting of SWI/SNF and P300 may trigger a more pronounced impact on SS18-SSX function.

**Figure 5:**
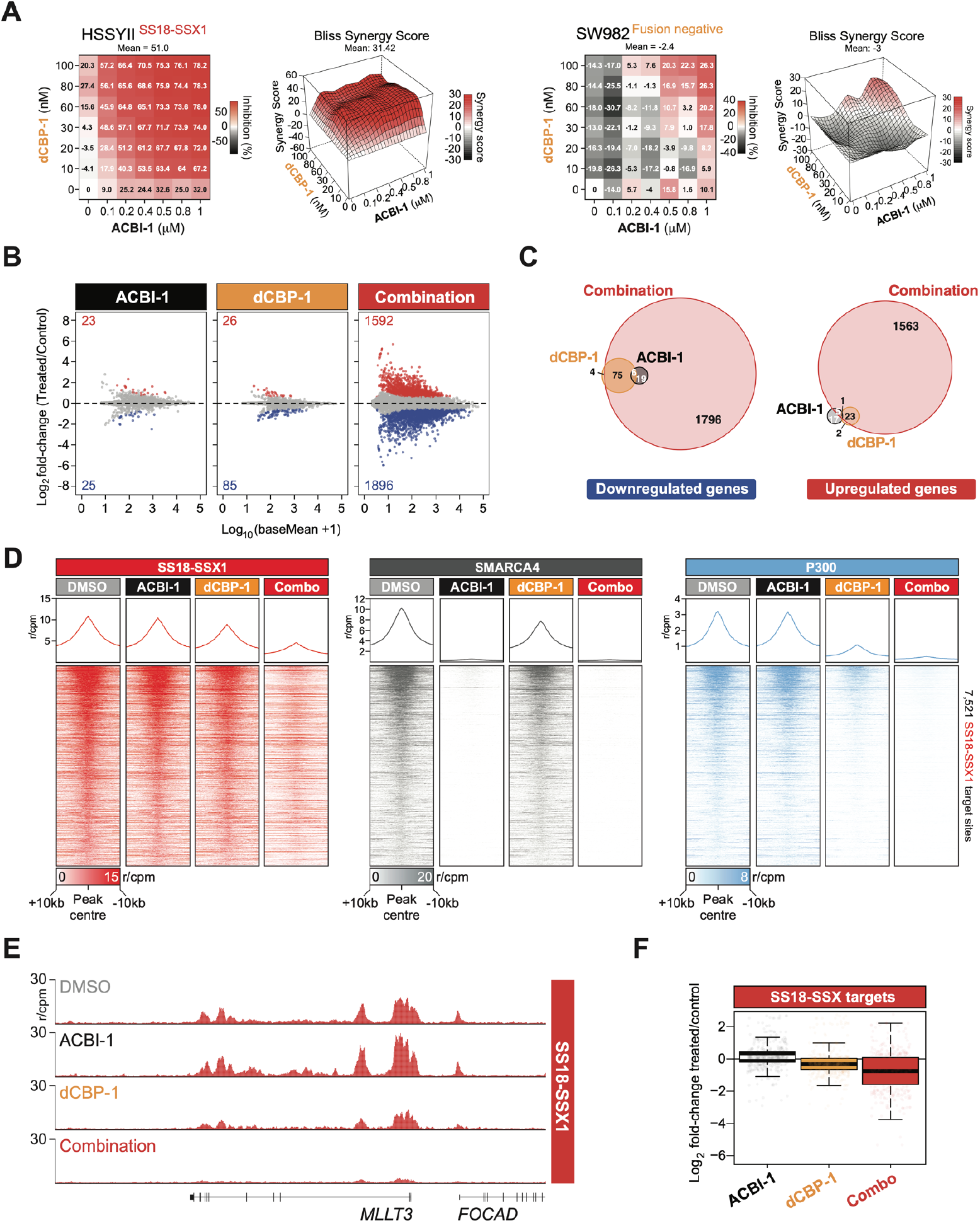
PROTAC targeting of SWI/SNF and P300 is highly synergistic in synovial sarcoma. **A**. Heatmaps showing percent viability response in the indicated cell lines in dose response matrices using dCBP-1 and ACBI-1 (left panels); corresponding synergy heatmaps and synergy contour plots (Bliss) of combination treatments with increasing synergy indicated in red (right panels). **B**. MA plots showing differential gene expression in HSSYII cells following 24hrs of single agent 400nM ACBI-1, 40nM dCBP-1 or combination treatment. Significantly differentially expressed genes are indicated in red (upregulated) and blue (downregulated). **C**. Venn diagrams showing the numbers and overlap of differentially expressed genes in single agent and combination treatments. **D**. Tornado and average plots showing the abundance of SS18-SSX, SMARCA4 and P300 in CUT&RUN-Rx of control (DMSO), 24hr single agent 400nM ACBI-1, 40nM dCBP-1 and combination treated HSSYII cells. Scale denotes reference-adjusted counts per million mapped reads (r/CPM). **E**. Genomic tracks showing SS18-SSX CUT&RUN-Rx signal at the indicated genomic loci in control (DMSO), 24hr single agent 400nM ACBI-1, 40nM dCBP-1 and combination treated HSSYII cells. **F**. oxplots showing Log2 fold-change in mRNA levels of SS18-SSX target genes in single agent and combination treated HSSYII cells.

Next, we performed global gene expression profiling to examine the transcriptional impact of combined degrader treatment. Low individual doses of ACBI-1 or dCBP-1 produced minimal transcriptional changes (**Figure 5B**), consistent with their limited phenotypic effects at these concentrations (**Figure 5A** and **S5A-B**). In striking contrast, the combination treatment elicited a robust transcriptional response, with more than 30-fold more differentially expressed genes (1,592 upregulated and 1,896 downregulated) than either single agent alone (**Figure 5B-C**). Consistent with the largely non-overlapping impacts of ACBI-1 and dCBP-1, their individual gene-expression signatures were largely distinct. However, both were encompassed within the combination responsive gene set, indicating that dual degradation broadly expands upon the transcriptional consequences of each compound (**Figure 5C**). KEGG pathway analysis revealed that DNA replication, cell-cycle, developmental, and cancer-associated pathways were among the most strongly downregulated terms, whereas apoptosis and autophagy were among the most upregulated (**Figure S5C**). Thus, combined degrader treatment profoundly remodels the transcriptional landscape and induces a strong phenotypic response in synovial sarcoma cells.

We next examined the impact of combined degradation on SS18-SSX chromatin localisation. At the low doses used in the combination setting, each degrader alone had only modest effects on fusion protein binding. Low-dose ACBI-1 treatment had little impact on overall SS18-SSX levels, whereas low-dose dCBP-1 caused a minor reduction (**Figure 5D**). In contrast, the combined treatment produced a significant genome-wide loss of SS18-SSX occupancy (**Figure 5D-E** and **S5D-E**), indicating that concurrent degradation of SMARCA2/4 and P300 synergistically impairs fusion recruitment to chromatin. Consistent with higher dose single agent treatments, low dose ACBI-1 displaced SS18-SSX preferentially from sites with low levels of P300, while dCBP-1 primarily impacted sites with high levels of P300 (**Figure S5F**). However, consistent with the marked synergy of these compounds in combination treated cells, SS18-SSX is profoundly reduced across all its target sites (**Figure S5F**). Finally, we examined how these changes impacted fusion target gene expression. Within the broad transcriptional response to combination treatment, we saw a preferential downregulation of direct SS18-SSX targets (**Figure S5G**). Consistent with the observed synergy, fusion targets were downregulated to a significantly greater degree compared to single-agent treatments (**Figure 5F**). Collectively, these data demonstrate combined degradation of SMARCA2/4 and P300/CBP profoundly disrupt SS18-SSX function eliciting strong therapeutic responses in sarcoma cells.

## DISCUSSION

Synovial sarcoma is driven by SS18-SSX, which has been assumed to rely on SWI/SNF chromatin remodelling activity for oncogenic transcription1. Our discoveries highlight that SS18-SSX drives disease associated gene expression independent of SWI/SNF complexes. Instead, the fusion depends on the acetyltransferase P300. These findings explain the limited efficacy of SWI/SNF targeting clinical trials in synovial sarcoma patients^23^. Importantly, they also provide a mechanistically anchored basis for implementing new therapeutic approaches for this challenging disease.

### The role of SWI/SNF complexes in synovial sarcoma

It has been well established that SS18-SSX incorporates into SWI/SNF complexes in sarcoma cells^12,13^. In doing so, it causes biochemical changes within complexes and leads to the redistribution of these formations on chromatin^13,19,20^. Given these discoveries, it is understandable how the prevailing notion, that altered SWI/SNF function is the central mechanistic aspect underlying disease development emerged. However, it is noteworthy that no studies have unequivocally demonstrated that SS18-SSX relies on SWI/SNF complex function to drive sarcoma development. Moreover, recent work has shown that SS18-SSX is still capable of driving sarcomagenesis in mouse models, even when SWI/SNF members including the cBAF components, Arid1a and Arid1b are deleted^48^. This supports the idea that the fusion is not fundamentally dependent on SWI/SNF activity to mediate its oncogenic effects. However, it is clear that the SWI/SNF complex is required to support binding of SS18-SSX to at least some of its target sites. Targeted degradation of SMARC2/4 and dissolution of SWI/SNF complexes leads to reduced levels of SS18-SSX at some target sites. The fact this does not broadly impact on fusion associated gene expression, supports that the fusion is not dependent on SWI/SNF activity for target gene expression. However, it does show that SS18-SSX needs to incorporate into intact SWI/SNF complexes to fully engage chromatin. The mechanistic aspects of this remain unclear. It could be that chromatin interaction interfaces within SWI/SNF contribute to binding and/or stabilisation of SS18-SSX containing complexes at target regions. Alternatively, it could be that interactions between intact SWI/SNF complexes and DNA-binding transcription factors or other chromatin regulators stabilise SS18-SSX binding. It will be important to continue efforts to discover the molecular aspects underlying chromatin engagement by SS18-SSX containing complexes to fully understand this disease.

### P300 specific dependency in synovial sarcoma

P300 and CBP have been pursued as therapeutic targets in several cancers, including some fusion protein driven diseases^49-52^. However, in many instances the function of these two proteins appears to be redundant. It was striking that our genetic screening experiments demonstrated that synovial sarcoma cells are uniquely dependent on P300 and not CBP. In the cell models examined in our study, the expression of each gene was comparable, indicating that this differential functional requirement is not simply related to differences in abundance. Instead, it implies that P300 executes some specific role(s) that integrate with SS18-SSX which CBP cannot accomplish. Given the general level of redundancy that exists between P300 and CBP, it is exciting that synovial sarcoma provides a context where there is a clear functional divergence. It will be important to pursue this in greater detail to begin understanding how and where these proteins functionally diverge. Discoveries in this regard will have important implications for understanding synovial sarcoma biology; and could highlight how to best exploit the therapeutic potential of P300 targeting. This specificity also raises the possibility that P300 selective small-molecules, could be used clinically to achieve patient responses with less systemic toxicity compared to dual P300/CBP targeting approaches. Related to this, despite the high levels of homology between P300 and CBP some drug development efforts are beginning to have success in attempts to develop P300 or CBP selective small-molecules^53,54^.

### Opportunities exploiting combination therapy

Aberrant chromatin regulatory fusion proteins, like SS18-SSX, are typically highly disease specific. However, the molecular mechanisms they exploit to promote disease associated gene expression often rely on general transcriptional co-factors1. The therapeutic targeting of such factors does present some challenges, since their activity is often required in many tissues and cell types. Certainly, the central importance of these disease driving fusion proteins and their associated co-factors, creates a therapeutic window wherein targeting such factors can provide relative therapeutic specificity. However, improvements in this regard would always be advantageous. Our discovery that co-targeting SWI/SNF and P300 provides significant therapeutic synergy is important in this regard. We were able to achieve profound viability impacts at significantly reduced drug doses in a combination setting. Something that was unique to SS18-SSX positive cells. This provides a rational basis for the design of future clinical testing exploiting this combination approach. Excitingly, there are clinical grade small-molecules targeting SMARCA2/4 and P300 already in testing or approved clinical use. Therefore, our discoveries could motivate the implementation of novel clinical testing in the near future.

## Acknowledgements

We are grateful to all members of the Brien lab who have contributed to the development of this work through ongoing input and scientific discussion. We are thankful to Duncan Sproul and Christine Rodger for sharing the TetO-GFP reporter cell lines. We also thank the University College Dublin (UCD) Genomics Core facility for assistance with NGS library sequencing, and the Institute of Genetics and Cancer Research Computing group for computational support. UCD mass spectrometry was supported by The Comprehensive Molecular Analytical Platform (CMAP) under The Science Foundation Ireland (SFI) Research Infrastructure Programme (18/RI/5702). Work in the Brien lab was supported by an SFI - Starting Investigator Research Grant (18/SIRG/5573); Worldwide Cancer Research (21-0271); and the UKRI (EP/X039633/1).

