## Supplemental information for "SS18-SSX co-opts P300 to sustain oncogenic transcription independent of SWI/SNF activity"

### Figure S1 - SWI/SNF chromatin dynamics in BRD9 PROTAC treated cells

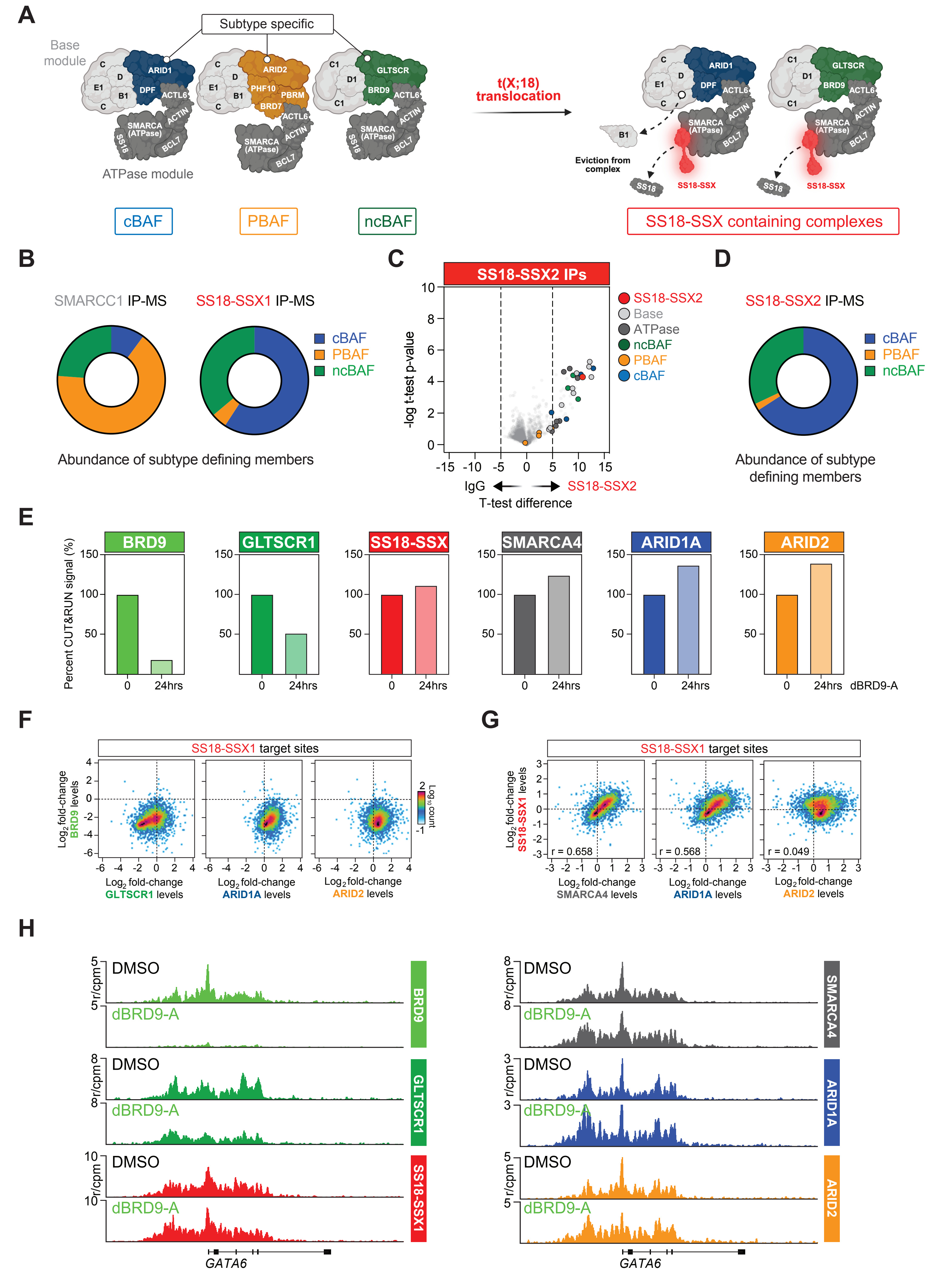

**Figure S1: SWI/SNF chromatin dynamics in BRD9 PROTAC treated cells**

**A.** Schematic representation of the SWI/SNF complex subclasses with ATPase and base modules, in addition to the subtype defining members indicated (left). Associated changes in cBAF and ncBAF formations following incorporation of SS18-SSX in synovial sarcoma cells are also shown (right).

**B.** Donut plot showing the proportions of signal derived from subclass defining SWI/SNF members in SMARCC1 and SS18-SSX purification mass spec experiments in HSSYII cells.

**C.** Volcano plot of SS18-SSX purification mass spec data from CME1 cells. SWI/SNF complex members are colour coded and indicated on the plot.

**D.** Donut plot showing the proportions signal derived from subclass defining SWI/SNF members in SS18-SSX purification mass spec experiments in CME1.

**E.** Bar plots showing percent total CUT&RUN-Rx signal at SS18-SSX bound regions in control and dBRD9-A treated HSSYII cells.

**F.** XY scatter plot correlating changes in BRD9 compared to GLTSCR1, ARID1A and ARID2 CUT&RUN-Rx signal at fusion protein target sites in 24hrs dBRD9-A 100nM treated HSSYII cells.

**G.** XY scatter plot correlating changes in SS18-SSX compared to SMARCA4, ARID1A and ARID2 CUT&RUN-Rx signal at fusion protein target sites in 24hrs dBRD9-A 100nM treated HSSYII cells.

**H.** Genomic tracks showing CUT&RUN-Rx signal of the indicated SWI/SNF members at the indicated genomic loci in control (DMSO) and 24hrs dBRD9-A 100nM treated HSSYII cells.

### Figure S2 - Targeting SMARCA2/4 in synovial sarcoma cells

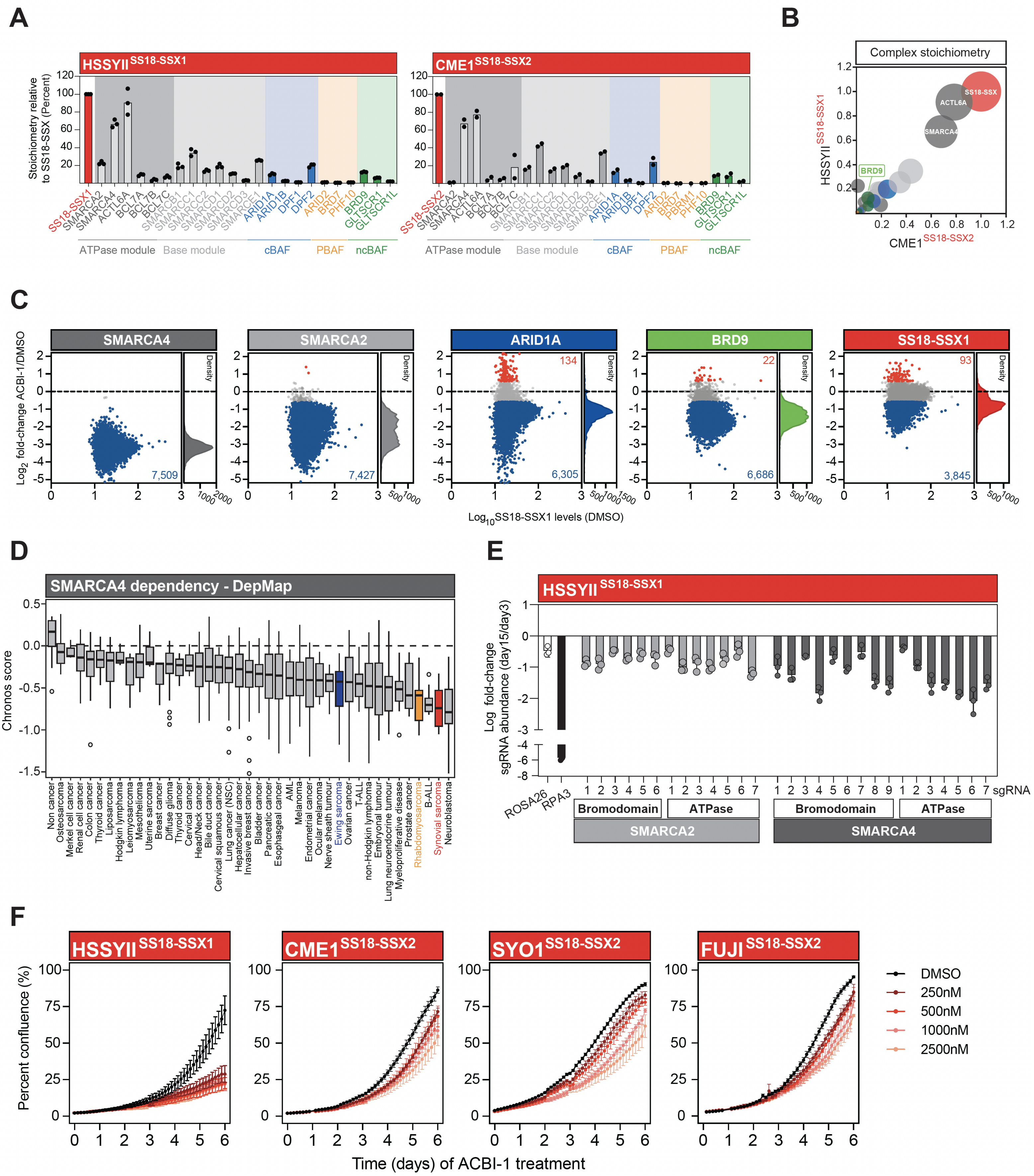

**Figure S2: Targeting SMARCA2/4 in synovial sarcoma cells**

**A.** Bar charts showing stoichiometry of each of the indicated SWI/SNF members in SS18-SSX purification mass spec experiments in HSSYII and CME1 cells.

**B.** Bubble plot showing stoichiometry of SWI/SNF members in SS18-SSX purifications in HSSYII and CME1 cells.

**C.** MA plots showing changes in SWI/SNF member CUT&RUN-Rx signal at SS18-SSX target sites following 24hrs of 1μM ACBI-1 treatment. Sites changing >1.5-fold are shown in red (increasing) or blue (decreasing) with the number of sites in these categories indicated.

**D.** Box plots showing DepMap Chronos score for SMARCA4 in cell lines of the indicated disease subtypes.

**E.** Growth competition assays in the indicated synovial sarcoma lines sgRNAs targeting the SMARCA2 or SMARCA4 bromodomain or ATPase domain (n = 3, data represents mean +/- SD).

**F.** Quantification of cell line growth in extended control (DMSO) and ACBI-1 treatments of the indicated synovial sarcoma cell lines.

### Figure S3 - P300 is functionally essential in synovial sarcoma cells

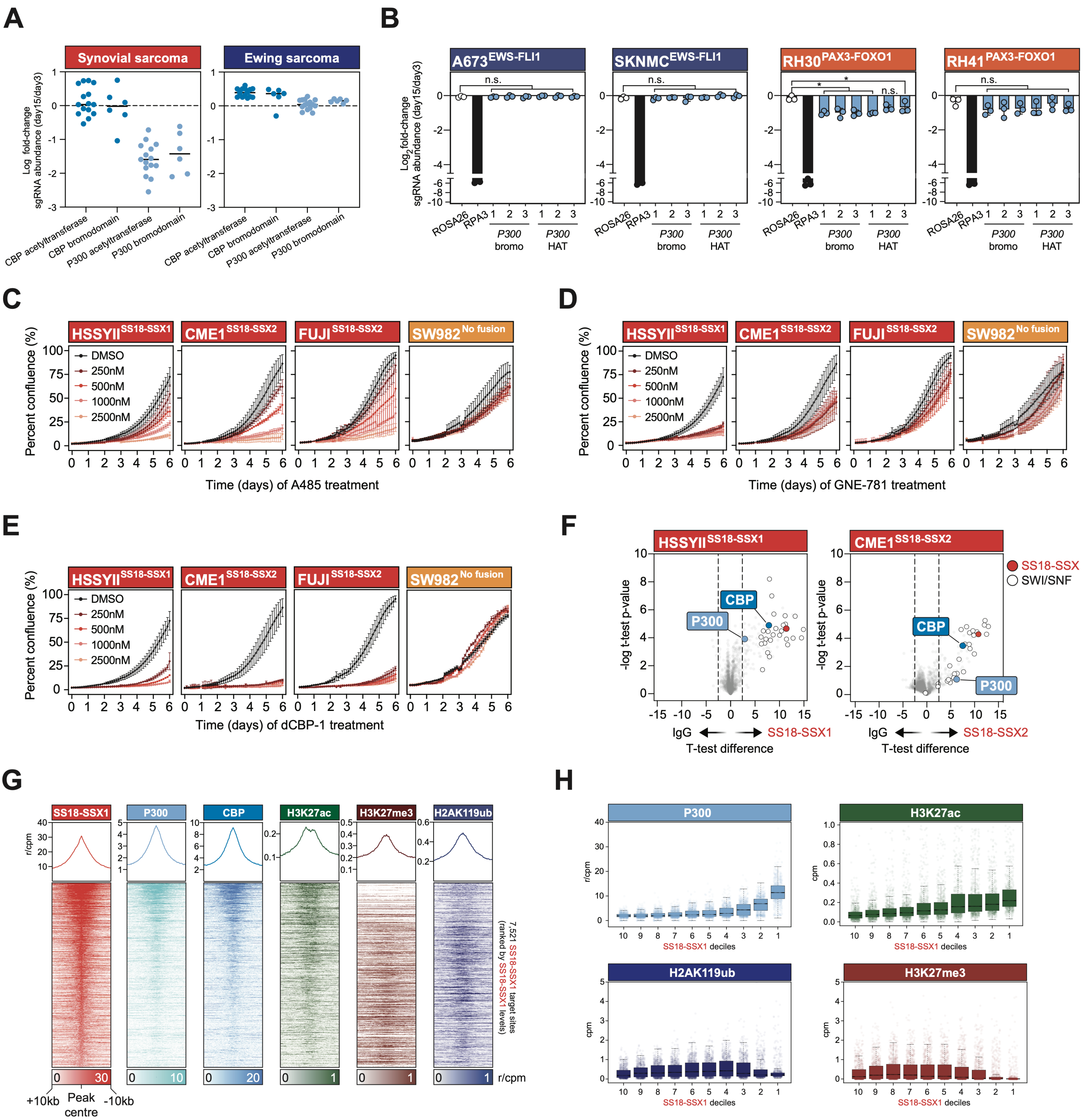

**Figure S3: P300 is functionally essential in synovial sarcoma cells**

**A.** Dot plot showing the drop out of individual sgRNAs targeting the CBP or P300 bromodomain or acetyltransferase domain in synovial and Ewing sarcoma cell lines. Each dot denotes the average change in sgRNA abundance in replicate experiments in multiple cell lines.

**B.** Growth competition assays in the indicated Ewings and rhabdomyosarcoma cell lines expressing P300 bromodomain or acetyltransferase targeting sgRNAs (n = 3, data represents mean +/- SD). P-values calculated using Student's t-test comparing against negative control sgRNA targeting ROSA26, * = P ≤ 0.05, ** = P ≤ 0.01.

**C.** Quantification of cell line growth in extended control (DMSO) and A-485 treatments of the indicated SS18-SSX positive and negative synovial sarcoma cell lines.

**D.** Quantification of cell line growth in extended control (DMSO) and GNE-781 treatments of the indicated SS18-SSX positive and negative synovial sarcoma cell lines.

**E.** Quantification of cell line growth in extended control (DMSO) and dCBP-1 treatments of the indicated SS18-SSX positive and negative synovial sarcoma cell lines.

**F**. Volcano plots of SS18-SSX purification mass spec data from HSSYII and CME1 cells. SWI/SNF complex are indicated on the plot, P300 and CBP are colour coded and indicated.

**G.** Tornado and average plots showing CUT&RUN signal for SS18-SSX, P300, CBP and the indicated histone modifications at SS18-SSX target regions in HSSYII cells.

**H** Boxplots showing CUT&RUN-Rx signal for P300 and the indicated histone modifications at SS18-SSX bound regions. Regions are ranked and sorted into deciles based on fusion protein abundance.

### Figure S4 - SS18-SSX binding dynamics in PROTAC treated cells

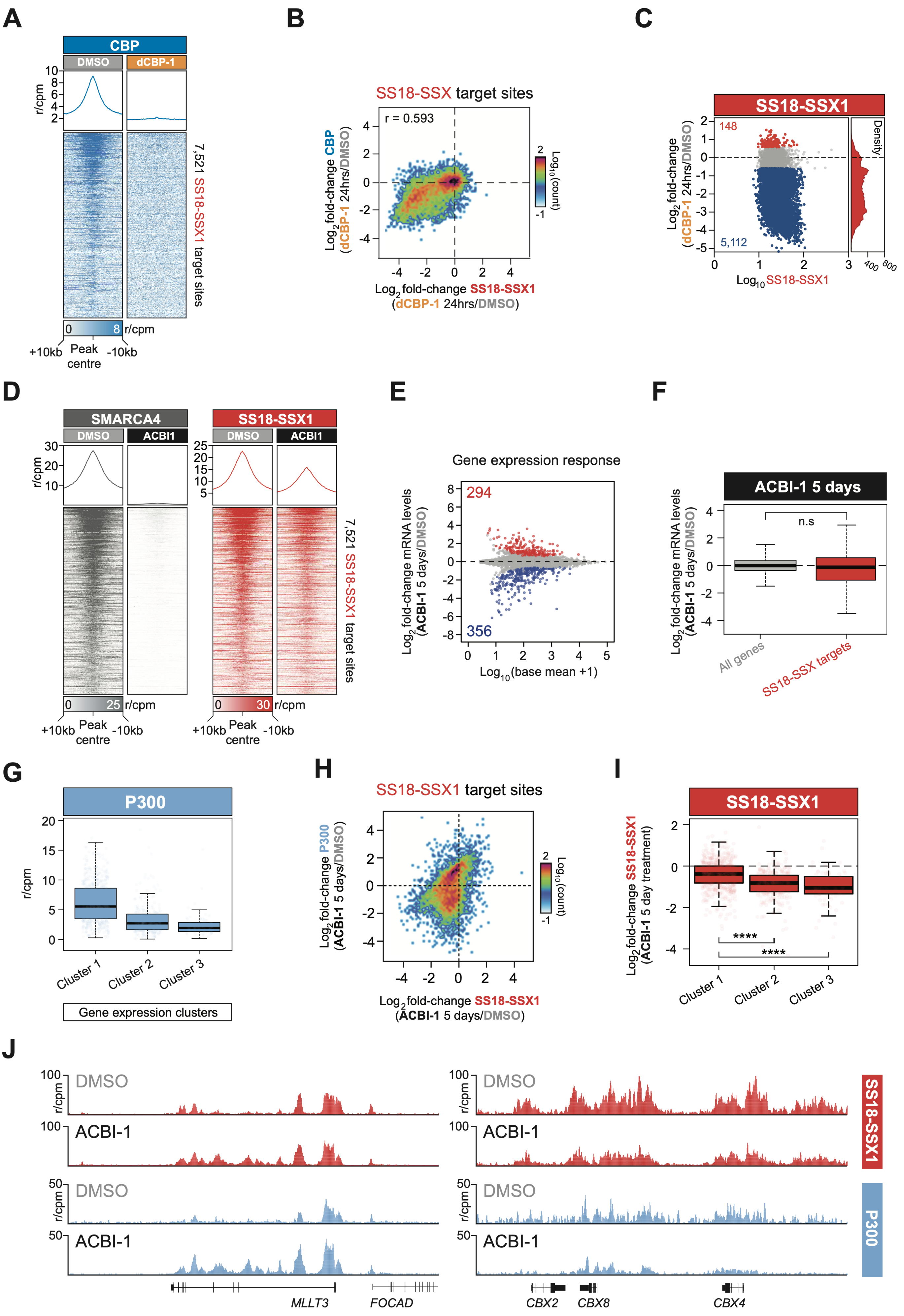

**Figure S4: SS18-SSX binding dynamics in PROTAC treated cells**

**A.** Tornado and average plots showing the abundance of CBP in CUT&RUN-Rx of control (DMSO) and dCBP-1 treated HSSYII cells. Scale denotes reference-adjusted counts per million mapped reads (r/CPM).

**B.** XY scatter plot correlating changes in CBP and SS18-SSX CUT&RUN-Rx signal at fusion protein target sites in 24hrs dCBP-1 500nM treated HSSYII cells.

**C.** MA plots showing changes in SS18-SSX CUT&RUN-Rx signal at fusion target sites following 24hrs of dCBP-1 500nM treatment. Sites changing >1.5-fold are shown in red (increasing) or blue (decreasing) with the number of sites in these categories indicated.

**D.** Tornado and average plots showing the abundance of SMARCA4 and SS18-SSX in CUT&RUN-Rx of control (DMSO) and 1μM ACBI-1 (5 days) treated HSSYII cells. Scale denotes reference-adjusted counts per million mapped reads (r/CPM).

**E.** MA plot showing differential gene expression in HSSYII cells following 5 days of 1μM ACBI-1 treatment. Significantly differentially expressed genes are indicated in red (upregulated) and blue (downregulated).

**F.** Boxplot showing $Log_2$ fold-change of direct SS18-SSX target genes (red) and all other genes (grey) in HSSYII cells following 5 days of 1μM ACBI-1 treatment.

**G.** Boxplot showing the abundance of P300 in CUT&RUN-Rx signal at clusters of differentially expressed direct fusion protein target genes (as per Figure 4G).

**H.** XY scatter plot correlating changes in P300 and SS18-SSX CUT&RUN-Rx signal at fusion protein target sites in 1μM ACBI-1 (5 days) treated HSSYII cells.

**I.** Boxplots showing $Log_2$ fold-change in SS18-SSX CUT&RUN-Rx signal at peaks associated with differentially expressed fusion target gene clusters (as per Figure 4G).

**J.** Genomic tracks showing SS18-SSX and P300 CUT&RUN-Rx signal at the indicated genomic loci in control (DMSO) and 5-day ACBI-1 treated HSSYII cells.

### Figure S5 - PROTAC treatment therapeutic synergy

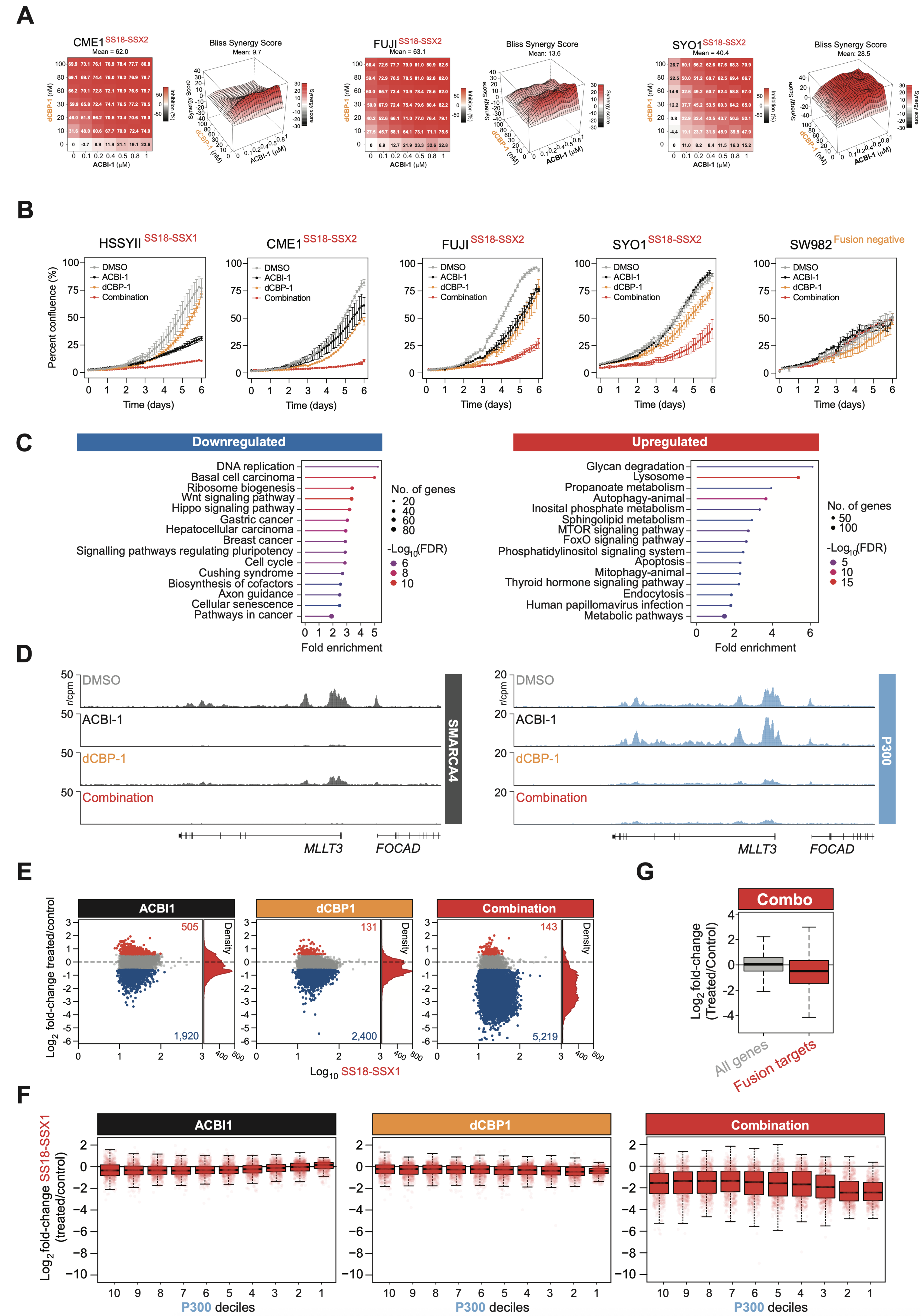

**Figure S5: PROTAC treatment therapeutic synergy**

**A.** Heatmaps showing percent viability response in the indicated cell lines in dose response matrices using dCBP-1 and ACBI-1 (left panels); corresponding synergy heatmaps and synergy contour plots (Bliss) of combination treatments with increasing synergy indicated in red (right panels).

**B.** Quantification of cell line growth in extended control (DMSO), single agent 400nM ACBI-1 or 40nM dCBP-1 and combination treatments of the indicated SS18-SSX positive and negative synovial sarcoma cell lines.

**C.** KEGG pathway analyses of genes significantly up and downregulated in combination 400nM ACBI-1 and 40nM dCBP-1 treatments (24hrs) in HSSYII cells.

**D.** Genomic tracks showing SMARCA4 and P300 CUT&RUN-Rx signal at the indicated genomic loci in control (DMSO), single agent 400nM ACBI-1, 40nM dCBP-1 and combination treated (24hrs) HSSYII cells.

**E.** MA plots showing changes in SS18-SSX CUT&RUN-Rx signal at fusion target sites following 24hrs of single agent 400nM ACBI-1, 40nM dCBP-1 or combination treatment. Sites changing >1.5-fold are shown in red (increasing) or blue (decreasing) with the number of sites in these categories indicated.

**F.** P300 decile boxplots showing $Log_2$ fold-change in SS18-SSX CUT&RUN-Rx signal at fusion protein target sites in single agent 400nM ACBI-1, 40nm dCBP-1 or combination treated (24hrs) HSSYII cells.

**G.** Boxplot showing $Log_2$ fold-change of direct SS18-SSX target genes (red) and all other genes (grey) in HSSYII cells following 24hrs of combination treatment.
