## Supplementary material for "SS18-SSX co-opts P300 to sustain oncogenic transcription independent of SWI/SNF activity": Materials and Methods

### Cell Culture and Lentiviral Production

All cell cultures were maintained in a humidified, normoxic incubator at 37ºC, 5% $CO_2$. HSSYII, SYO1, SW982, HEK293T C2C12 cells were cultured in DMEM media (Sigma) supplemented with 10% foetal bovine serum and 100 U/ml Penicillin/100 µg/ml Streptomycin (Gibco). CME1 and FUJI cells were cultured in RPMI media (Sigma) supplemented with 10% foetal bovine serum and 100 U/ml Penicillin/100 µg/ml Streptomycin (Gibco). Mouse embryonic stem cells were cultured under 2i/LIF conditions in GMEM (Sigma) supplemented with 20% heat-inactivated FBS, penicillin/streptomycin (100 µg/mL), 50 µM β-mercaptoethanol, GlutaMAX (1:100), and plated on 0.1% gelatin coated dishes.

Lentiviral particles were generated by co-transfection of HEK293T cells with a lentiviral expression vector or pooled sgRNA library with a viral packaging (PAX2) and envelope (VSV-G) vectors using PEI in accordance with standard protocols. Viral supernatants were collected between 24-72hrs post-transfection and either used directly for infection of target cells after filtering through a 0.45µm syringe filter and addition of 8.5 mg/ml Polybrene; or concentrated by ultracentrifugation (20,000 rpm, 2hrs) before being resuspended in 0.5-1ml of culture media and used for infection of target cells.

### Endogenous protein immunoprecipitation and mass spectrometry

Harvested cells were washed twice with cold PBS and washed pellets resuspended with Buffer A (25mM HEPES pH 7.6, 5mM $MgCl_2$, 25mM KCl, 0.05mM EDTA, 10% (v/v) glycerol, 0.1% (v/v) NP40, 1mM DTT and 1mM PMSF) and kept on ice for 10 minutes. Samples were centrifuged at 1,500rpm for 10 minutes at 4ºC and the supernatant was discarded. To lyse nuclei, pellets were resuspended in Buffer C (10mM HEPES pH 7.6, 3mM $MgCl_2$, 100mM KCl, 0.5mM EDTA, 10% (v/v) glycerol, 1mM DTT and 1mM PMSF). Next, 10% (v/v) 3M $(NH_4)_2SO_4$ prepared in Buffer C was added to the samples, which were rotated at 4ºC for 20 minutes. Samples were ultracentrifuged at 350,000 rcf for 15 minutes at 4ºC in a Beckman Coulter Optima L-100XP using a SW 55Ti rotor. The supernatant was collected and 300mg of $(NH_4)_2SO_4$ was added for each 1mL of sample, to precipitate nuclear extracts. Samples were vortexed and kept on ice for 15 minutes before being ultracentrifuged using the same parameters as above. Pellets were resuspended in IP buffer (300mM NaCl, 50mM Tris-HCl pH 7.5, 1mM EDTA, 1% (v/v) Triton-X-100, 1mM DTT and 1mM PMSF) and Bradford assays were performed to determine protein concentration. Benzonase Nuclease (Sigma) and 1µg of the respective antibody were added to each IP and samples were rotated at 4ºC overnight. the following day Protein G Dynabeads (Thermo Fisher) were added to each sample, followed by rotation at 4ºC for 2.5 hours. Beads were washed 5x with IP buffer, before being processed for immunoblotting or mass spectrometry sample preparation.

In-solution tryptic digestions were performed as described previously[55]**.** Samples were run on a Bruker timsTof Pro mass spectrometer connected to an Evosep One liquid chromatography system. Tryptic peptides were resuspended in 0.1% formic acid and each sample was loaded on to an Evosep tip and separated on a Evosep EV1137 Performance Column – 15 cm x 150 um, 1.5 um. The Evosep tips were placed in position on the Evosep One, in a 96-tip box. The autosampler is configured to pick up each tip, elute and separate the peptides using a set chromatography method (15 samples a day)[56]. The mass spectrometer was operated in positive ion mode with a capillary voltage of 1650 V, dry gas flow of 3 l/min and a dry temperature of 180 °C. All data was acquired with the instrument operating in trapped ion mobility spectrometry (TIMS) mode. Trapped ions were selected for ms/ms using parallel accumulation serial fragmentation (PASEF). A scan range of (100-1700 m/z) was performed at a rate of 5 PASEF MS/MS frames to 1 MS scan with a cycle time of 1.03s[57].

**CUT&RUN-Rx**

Cells were harvested using Accutase (STEMCELL Technologies) and counted. Following counting, 100,000 mouse ESCs or C2C12 cells were spiked-in to 900,000 synovial sarcoma cell suspensions (10% spike-in). Cells were fixed in culture media containing 0.1% formaldehyde at room temperature for 1 min. Formaldehyde was quenched with glycine at 0.125 M, followed by a 5-min incubation at room temperature. Fixed cells were washed with PBS. Fixed cells were incubated on ice for 10 min in nuclear extraction buffer (20mM HEPES pH 7.5, 10mM KCL, 0.1% Triton X-100, 20% Glycerol,1X protease inhibitor cocktail and 0.5mM spermidine). Extracted nuclei were collected by centrifugation (4°C, 600g) and resuspended in cold nuclei extraction buffer (100$\mu$L per sample). CUT&RUN was performed following Epicypher CUT&RUN Protocol v1.5.1. CUT&RUN DNA was purified using the Monarch PCR & DNA Cleanup Kit (New England Biolabs) in accordance with the manufacturer's instructions.

**CUT&RUN-Rx Library Preparation**

Purified DNA was quantified using a Qubit fluorimeter (Invitrogen). 1-10 ng of DNA was used to generate CUT&RUN-Rx libraries with the NEBNext Ultra II DNA Library Prep Kit for Illumina (New England Biolabs) as per the manufacturer's instructions. Library DNA was quantified using the Qubit, and size distributions were ascertained on a TapeStation (Agilent) using the D1000 ScreenTape assay reagents (Agilent; 5067- 5583). This information was used to calculate pooling ratios for multiplex library sequencing. Pooled libraries were diluted and processed for 38-bp paired-end sequencing on an Illumina NextSeq500 instrument using the NextSeq 500 High Output v2 kit (Illumina; FC-404-2005) in accordance with the manufacturer's instructions. Or alternatively, sequenced commercially (Novogene) using 300-bp paired-end sequencing on the NovaSeq system.

**CUT&RUN sequencing data analysis**

A hybrid human–mouse reference genome was generated by appending hg38 to mm10 (with an mm10 prefix added to chromosome names) and indexed with bowtie2. Reads were aligned using bowtie2 in --very-sensitive mode. Using SAMtools, reads with MAPQ < 2 were removed, human and mouse reads were separated, duplicates were removed, and SAM files were converted to BAM. Spike-in normalization factors were calculated using the reference-adjusted reads per million (RRPM) method. RRPM-normalized bigWig files were generated with deepTools bamCoverage using the appropriate scale factor. Peaks were called with macs2 (paired-end mode) using matched IgG controls and an fdr <0.05. BroadPeak outputs were converted to BED format and filtered using bedtools intersect to remove regions overlapping a custom hg38 blacklist. Peaks were annotated with HOMER. Consensus peaks were defined across replicates using MSPC (parameters: -r Bio -w 1e-4 -s 1e-8). Average RRPM bigWigs were generated using BigWigAverage.

**Quant-seq sample prep and sequencing**

Total RNA was isolated from control and compound treated synovial sarcoma cell cultures using the RNeasy kit (Qiagen). The quality of extracted RNA was confirmed using the TapeStation (Agilent) with the RNA ScreenTape assay reagents (Agilent; 5067-5576). Total RNA (500 ng) was used as input material from each sample for library preparation. Libraries were generated using the QuantSeq 3′ mRNA-Seq Library Prep Kit FWD for Illumina (Lexogen; 015.24) in accordance with the manufacturer's instructions. Library DNA was quantified using the Qubit, and size distributions were ascertained on a TapeStation (Agilent) using the D1000 ScreenTape assay reagents (Agilent; 5067-5583). This information was used to calculate pooling ratios for multiplex library sequencing. Pooled libraries were diluted and processed for 75-bp single-end sequencing on an Illumina NextSeq instrument using the NextSeq 500 High Output v2 kit (75 cycles) (Illumina; FC-404-2005) in accordance with the manufacturer's instructions. Or alternatively, sequenced commercially (Novogene) using 300-bp paired-end sequencing on the NovaSeq system

**Quant-seq data analysis**

Only the forward read from the paired-end libraries was used for downstream processing. Adapter trimming and quality filtering were performed with the BBDuk module from the BBTools suite, following the manufacturer's recommendations. High-quality reads were aligned to the human genome (hg38) using the STAR aligner. Gene-level read counts were generated with **htseq-count**, and differential expression analysis was carried out using the **DESeq2** package in R, applying a threshold of >1.5-fold change and a Benjamini–Hochberg adjusted $P$ value < 0.05.

**Fusion protein transcriptional reporter assays**

The TetO-GFP reporter cell line was a kind gift from Dr. Duncan Sproul and Dr. Christine Rodger and was generated using a Bxb1-recombinase genomic landing pad system in HEK293T cells. Into this background we introduced an SS18-SSX fusion protein with an N-terminal TetR-DNA binding domain. Expression of the GFP reporter locus was measured using the Guava easyCyte Flow Cytometer (Cytek). Reporter cells were treated with the indicated compound doses and the impact of these treatments on reporter gene expression was measured by flow cytometry.

**Pooled CRISPR screening**

The human epigenetic domain U6-sgRNA-EFS-GFP targeting library was pooled at equimolar ratio and used to generate lentiviral supernatant as described above. The total number of synovial and Ewing's sarcoma target cells for infection was chosen to achieve at least 500-fold representation of each sgRNA in the cell population. To ensure that a single sgRNA was transduced per cell, the viral volume for infection was chosen to achieve an MOI of 0.3–0.4. Genomic DNA was extracted time points the QiAamp DNA mini kit (Qiagen #51304), following the manufacturer's instructions. To maintain >500X sgRNA library representation, 16–20 independent PCR reactions were used to amplify the sgRNA cassette, which were amplified for 20 cycles with 100–200 ng of starting gDNA using the 2x Phusion Mix (Thermo Scientific #F-548). The PCR products were pooled and end repaired with T4 DNA polymerase (NEB), DNA polymerase I (NEB), and T4 polynucleotide kinase (NEB). A dATP overhang was added to the end-repaired DNA using Klenow DNA Pol Exo- (NEB). The DNA fragment was then ligated with diversity-increased barcoded Illumina adaptors followed by five pre-capture PCR cycles. Barcoded libraries were pooled at equal molar ratio and subjected to massively parallel sequencing using a Mi-Seq instrument (Illumina) using paired-end 150 bp reads (MiSeq Reagent Kit v2; Illumina MS-102–2002). The sequence data were trimmed to contain only the sgRNA sequence then mapped to the reference sgRNA library without allowing any mismatches. The read counts were then calculated for each individual sgRNA. To compare the differential representation of individual sgRNAs between day 3 and day 15 time points, the read counts for each sgRNA were normalized to the counts of the negative control ROSA26 sgRNA.

**Negative selection assays**

Cas9-expressing synovial, Ewing or rhabdomyosarcoma cell lines were transduced with sgRNA-GFP expressing lentivirus at a low MOI and passaged without selection. The percentage of GFP positive cells was measured at each passage using the Guava easyCyte Flow Cytometer (Cytek). The relative proportion of GFP positive cells was monitored throughout this serial cell culture.

**Compound dose response assays**

For dose response viability assays, cells were plated in 96-well tissue culture plates (5000 cells/well) in media containing DMSO or the desired concentration of each compound. After 72hrs of treatment, CellTitreGlo (Promega) was used to determine ATP-dependent luminescence as an approximation of cell viability.

**Live Cell Imaging**

Cells were plated in 384-well format and treated with compounds at the indicated doses. 384-well plates were loaded into an Incucyte SX5 live cell optical module (Sartorius). Using the phase channel (10x objective), images were taken every 3hrs for a total of 144hrs. Quantification of cell confluence over time from these phase images was carried out using the built-in Incucyte analysis module (segmentation adjustment 0.1-0.3, Classic Confluence mode, minimum area filter 200 $\mu m^2$).

**Drug synergy assays**

For synergy assays cells (1000 cells/well) were seeded in 384-well plates. After ~20 hours, cells were dosed at varied concentrations or each compound, using a D300e Digital Dispenser (Tecan), as 7x7 combination matrices (n=3) with multiple DMSO only controls. All wells were standardised to 0.1% (v/v) DMSO and incubated for a further 72 hours before cell viability quantification using CellTiterGlo (Promega)

CellTiterGlo outputs for each matrix were normalised to DMSO wells as percent cell growth inhibition. The normalised data (n=3) was uploaded to SynergyFinderPlus (synergyfinder.org) for analysis and interpretation of potential synergistic combinations, using multiple mathematical models (Bliss, Loewe, Zero Interaction Potency (ZIP) & High Specific Activity (HSA)).
